# Real-time analysis of nanopore-based metagenomic sequencing from orthopaedic device infection

**DOI:** 10.1101/220616

**Authors:** Nicholas D Sanderson, Teresa L Street, Dona Foster, Jeremy Swann, Bridget L. Atkins, Andrew J. Brent, Martin A. McNally, Sarah Oakley, Adrian Taylor, Tim E A Peto, Derrick Crook, David W Eyre

## Abstract

Prosthetic joint infections are clinically difficult to diagnose and treat. Previously, we demonstrated metagenomic sequencing on an Illumina MiSeq replicates the findings of current gold standard microbiological diagnostic techniques. Nanopore sequencing offers advantages in speed of detection over MiSeq. Here, we compare direct-from-clinical-sample metagenomic Illumina sequencing with Nanopore sequencing, and report a real-time analytical pathway for Nanopore sequence data, designed for detecting bacterial composition of prosthetic joint infections.

DNA was extracted from the sonication fluids of seven explanted orthopaedic devices, and additionally from two culture negative controls, and was sequenced on the Oxford Nanopore Technologies MinION platform. A specific analysis pipeline was assembled to overcome the challenges of identifying the true infecting pathogen, given high levels of host contamination and unavoidable background lab and kit contamination.

The majority of DNA classified (>90%) was host contamination and discarded. Using negative control filtering thresholds, the species identified corresponded with both routine microbiological diagnosis and MiSeq results. By analysing sequences in real time, causes of infection were robustly detected within minutes from initiation of sequencing.

We demonstrate initial proof of concept that metagenomic MinION sequencing can provide rapid, accurate diagnosis for prosthetic joint infections. We demonstrate a novel, scalable pipeline for real-time analysis of MinION sequence data. The high proportion of human DNA in extracts prevents full genome analysis from complete coverage, and methods to reduce this could increase genome depth and allow antimicrobial resistance profiling.

## Background

Joint replacement surgery may be complicated by prosthetic joint infections (PJI), a rare but serious event occurring up to five or more years post-operatively[1]. Recent studies in England of joint revisions undertaken for infection report an increase in prevalence for both knee and hip revisions [2, 3]. Improvements in speed and accuracy of diagnosis may improve outcomes following revision surgery by allowing more targeted therapy. PJI diagnosis can be challenging as infections may be associated with biofilms that colonise the orthopaedic devices [4], caused by fastidious or slow-growing organisms that are not detectable by culture or from patients who have received prior antibiotics. Although culture of multiple periprosthetic tissue (PPT) samples remains the gold standard for microbial detection, it is relatively insensitive, with only approximately 65% of causative bacteria detected even when multiple PPT samples are collected [5–7].

Development of molecular methods, such as multiplex-PCR, can be more sensitive in detection of PJI but are restricted by choice of primers to detect specific bacteria. An alternative is the use of metagenomic shotgun sequencing that can detect all bacteria directly from a sample. Sequencing directly from samples can provide accurate diagnostic information for PJIs when compared to laboratory culture and can also detect additional organisms[8, 9].

Using third generation sequencing technology, developed by Oxford Nanopore Technologies (ONT) and Pacific Biosciences (PacBio), longer read lengths in faster turnarounds are possible. The ONT MinlON potentially could allow analysis to be conducted in real-time with obvious advantages to clinical diagnosis of infection. Examples of metagenomic pathogen studies using MinlON include viral detection from serum [10] and bacteria from urines and pleural effusions [11, 12]. These previous studies have shown proof-of-principle for direct from sample clinical sequencing using ONT MinlON. However, PJI sequencing has a further challenge of high human DNA contamination which require specific laboratory preparation and bioinformatic analyses to overcome.

Here we describe proof-of-principle for the use of ONT MinION sequencing for the diagnosis of PJI when compared to standard microbiological culture and Illumina sequencing. We describe an analysis work-flow that differentiates between predicted infection species and background contamination and can be run during sequencing for real-time species detection.

## Methods

### Samples

Samples used in this study were collected by the Bone Infection Unit at the Nuffield Orthopaedic Centre (NOC) in Oxford University Hospitals (OUH), UK, as previously described [8]. 9 samples previously assessed by Illumina MiSeq sequencing were chosen for further analysis by ONT MinlON sequencing. Samples were chosen from the remaining DNA extracts that had sufficient DNA to either be sequenced directly, or amplified and sequenced, and to represent a range of disparate species and compositions.

### DNA preparation and sequencing

Libraries were prepared for sequencing on an Oxford Nanopore MinION (Oxford Nanopore Technologies (ONT)) using genomic DNA previously extracted from sonication fluid samples [8]. Samples 259, 312, 335, 352 and 354 were prepared using the 1D genomic DNA by ligation protocol (SQK-LSK108) (ONT). Samples 229, 249, 506 and 509 had insufficient DNA for this protocol so were prepared using either a PCR-based protocol for low input genomic DNA with modified primers (DP006_revB_14Aug2015), followed by rapid sequencing adapter ligation (ONT) (sample 229) or the ID low input genomic DNA with PCR protocol (SQK-LSK108) (ONT) (samples 249, 506 and 509). Briefly, the protocols comprise DNA end-repair and dA-tailing (NEBNext Ultra II End Repair/dA-Tailing Module, New England Biolabs (NEB), Ipswich, MA, USA) followed by purification using AMPure XP solid phase reversible immobilisation (SPRI) beads (Beckman Coulter, High Wycombe, UK); Sequencing adapter ligation (Blunt/TA Ligase Master Mix, NEB) followed by additional SPRI bead purification. For the samples with insufficient DNA requiring PCR amplification, additional steps between end-repair and sequencing adapter ligation included; PCR adapter ligation (Blunt/TA Ligase Master Mix, NEB) followed by SPRI bead purification; PCR amplification (Phusion High Fidelity PCR Master Mix, NEB) with 18 cycles (samples 229 and 249) or 24 cycles (samples 506 and 509) followed by additional SPRI bead purification. Samples were sequenced on FLO-MIN105 (v.R9) (sample 229) or FLO-MIN106 (v.R9.4) (all other samples) SpotON flowcells.

### PCR analysis of sample 354a

Quantitative real-time PCR (q-PCR) was performed for sample 354a to determine relative amounts of both *Arcanobacterium haemolyticum* and *Fusobacterium nucleatum* DNA in the original sonication fluid genomic DNA extract. qPCR was performed on a Stratagene MX3005P QPCR System (Agilent Technologies, Santa Clara, CA, USA) using Luna Universal Probe qPCR Master Mix (New England Biolabs, Ipswich, MA, USA). For *A. haemolyticum,* primers and probe were designed to target the phospholipase D gene: forward primer ATGTACGACGATGAAGACGCG (previously published, [13]), reverse primer TTGATTGCGTCATCGACACT, probe [6FAM]-TTGGTAGTGCGGCTGCTGCGCC-[TAM]. For *F. nucleatum,* primers and probe were designed to target the *nusG* gene: forward primer CAGCAACTTGTCCTTCTTGATCC, reverse primer CTGGATTTGTAGGAGTTGGTTC, probe [6FAM]-AGACCCTATTCCTATGGAAGAGGAAGAAGTA-[TAM]. Reactions were performed in 20μl with 2ul of template DNA, 0.4μM of each primer and 0.2μM of the probe. Cycling conditions were an initial denaturation at 95°C for 1 minute, followed by 40 cycles of 95°C denaturation for 15 seconds and 60°C extension for 30 seconds. Genomic DNA, extracted from cultures of *A. haemolyticum* (Type Strain NCTC 8452) and *F. nucleatum subspecies vincentii* (Type Strain ATCC 49256), was diluted to 100,000 genome copies per μl then serially diluted to 10 genome copies per μl and used to create copy number standard curves for both species. Negative controls, replacing template DNA with water, were also performed. All reactions were performed in triplicate.

### Bioinformatics analysis

We assembled an analysis pipeline for detection of bacterial pathogens using ONT MinION sequencing of orthopaedic device infections. The pipeline includes filtering steps for the genetic sequence data that have been tuned on seven positive samples with known infections and two culture negative samples.

The analysis was performed within a Nextflow workflow [14] with the software contained within a Singularity [15] image generated from a Docker repository [16]. This workflow and software are available for public use, [17], with our intention for the analysis to be reproducible or replicable with other datasets on most systems.

The workflow, CRuMPIT, has three major components, as shown in Figure 1. The first monitors the output of a MinION device or devices and creates batches of fast5 files (default 1000) as they are written to a storage drive location, Figure 1 (a,b). The second receives the fast5 files and uses a Nextflow workflow that basecalls data to be classified and aligns them to specific reference sequences with results pushed to a database, Figure 1 (c). Thirdly, analysis results including species identified, are determined and continually updated as the run progresses, Figure 1 (d).

**Figure 1.**
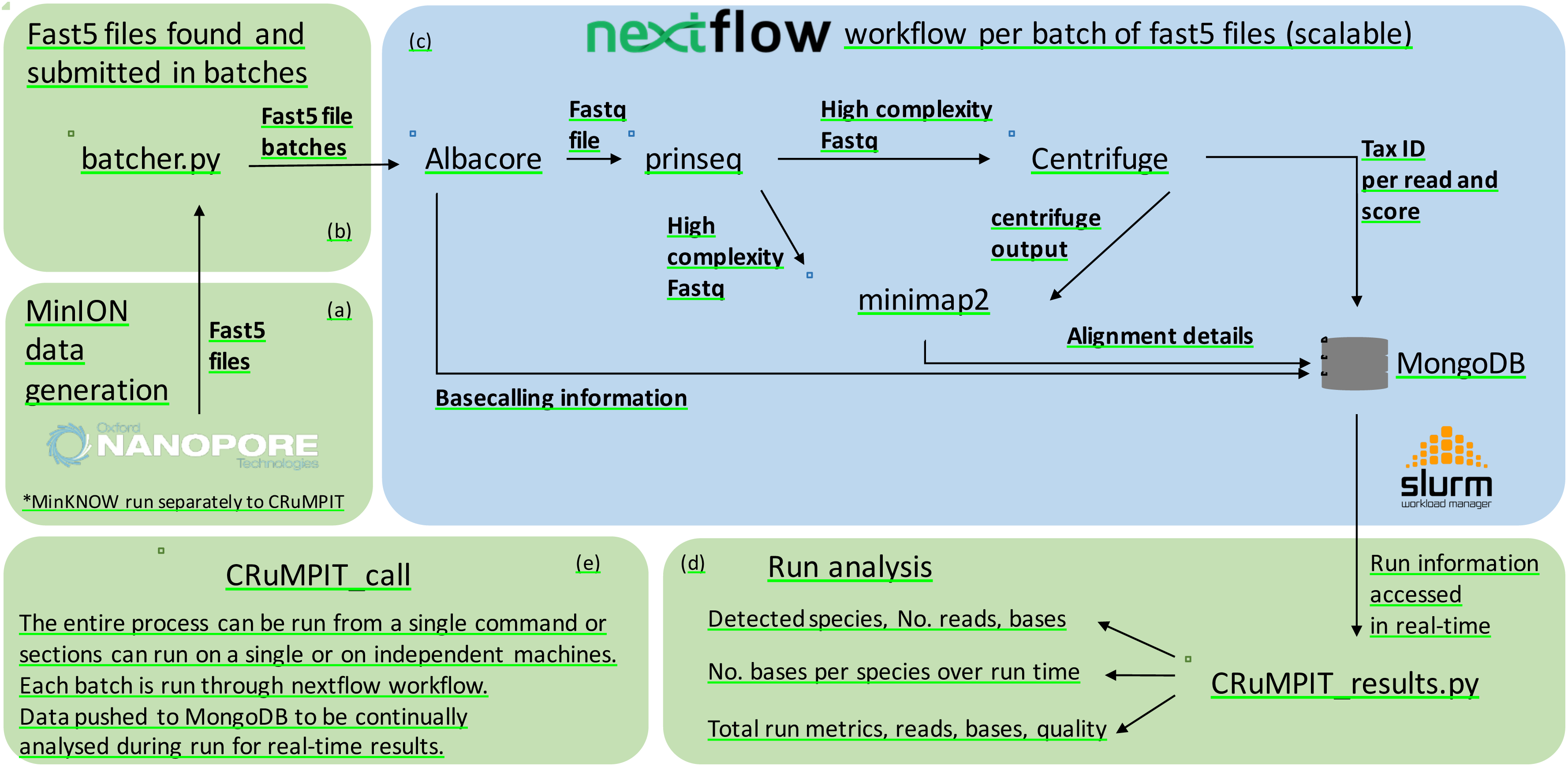
Diagram of analysis process. (a) MinION sequencing using MinKNOW (runs outside of CRuMPIT). (b) Fast5 files are detected and submitted as batches for the Nextflow workflow. (c) Nextflow workflow which is contained within a singularity image and can be distributed across a cluster (SLURM used here) or on a local machine. (d) Run analysis using data pushed to a MongoDB database, this can be conducted separately on any machine with network access to the database. Each component (green or blue rounded rectangle) of CRuMPIT can be run independently from the same or different networked computers, (e) or the entire process can be run from a single program. Square rectangles represent programs, some of which are within python wrappers. Arrows represent direction of data transfer within the workflow or between componants.

During the progression of this project, ONT have released several different software applications for basecalling, with each version improving accuracy [18]; we used the most up to date and reliable version at the time of sequencing. Basecalling from the fast5 files used different versions of either Metrichor (dragonet), MinKNOW-Live or ONT Albacore, Table 3. Fastq files were generated from the Metrichor or MinKNOW basecalled fast5 files using fast5watcher.py (commit b88e14a)[19] for downstream analysis. Albacore is now used as the basecaller within the CRuMPIT workflow, with sequences basecalled directly to fastq files for analysis.

To minimise spurious read classifications caused by repeat regions, sequences within the fastq files were separated based on molecular complexity, with only high complexity reads analysed further. Complexity was calculated using a dust score threshold of seven with prinseq-lite-0.20.4[20] which removes reads containing sequences consisting only of homopolymer, dipolymer and triploymer repeats.

Centrifuge [21] was used to classify sequencing reads to a taxomic identifier. We used Centrifuge instead of Kraken[22] for this analysis because the initial starting match uses kmers of length 16, which is more suited to the Nanopore error profile compared to Kraken where databases are built with a default kmer size of 31. Additionally, the Centrifuge indexes require significantly less storage and memory compared to Kraken. A Centrifuge index [21] was constructed using bacterial and viral genomes downloaded from NCBI RefSeq as of 03-March-2017, and the human reference genome (GRCh38). Low complexity regions with a dust score greater than 20 in the reference sequences were masked using dustmasker (v 1.0.0, NCBI). Alternatively, the precompiled “P_compressed_b+v+h” available to download from the Centrifuge authors was also used, yielding very similar results to our database. We used our database for this analysis because it is a more recent and complete dataset. However, for ease of reproducibility, the precompiled databases can also be used.

Sequences with a taxonomic id, or a descendant, that belonged to a list of bacterial reference genome sequences downloaded from NCBI RefSeq, were mapped using minimap2 [23] (v2.2-r409). To be considered for detection, bacterial species were first classified by Centrifuge with a score of 150 or greater with over 10% of the classified bacterial bases. The score of 150 was chosen as a suitable cutoff after several thresholds were tested, Supplemental figure 1. To remove spurious hits and background lab contamination, species were reported if they accounted for over 10% of the classified bacterial bases by Centrifuge which also removed the majority of negative control hits, Supplemental figure 2. Alternatively, a read number threshold could have been chosen, however the margin of proportional read numbers was deemed too narrow between positive samples and negative controls. Therefore, a further mapping step was added to validate the Centrifuge classification.

**Figure 2.**
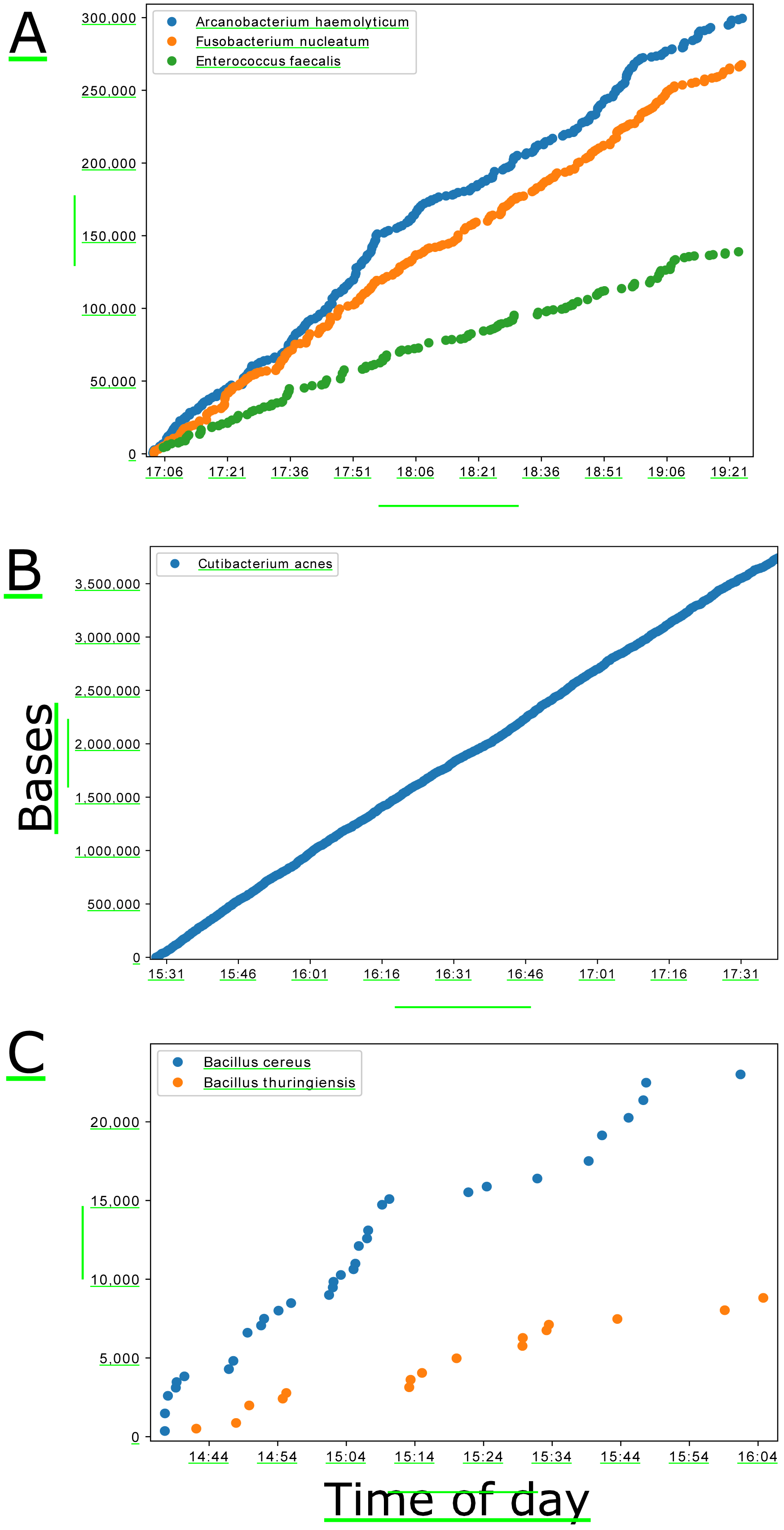
Cumulative bases classified by Centrifuge and minimap2 reference alignment over the first few hours of sequencing on the MinlON. Each marker on the plots represents a new sequence classified. Times are on the day of sequencing and taken from the read timestamp and doesn’t include bioinformatic time. Three samples shown showcasing the best and worst performers. (A) Sample 354a containing three different species. (B) Sample 249a containing Cutibacterium acne. (C) Sample 352a containing two different Bacillus species.

To be confirmed as a positive the mapped reads required a mapping quality score (mapq) of 50 or above and had to account for greater than 1% of the classified bacterial bases. Mapq 50 was used to ensure high quality alignments and helped to remove any remaining indiscriminate alignments, Supplemental figure 3. The 1% bases threshold was used after plotting bases over reads for positive samples and negative controls, Supplemental figure 4. However, if a detection species meets these criteria, the mapped reads can have any Centrifuge score and are included in further analysis. Therefore, more reads can be included if mapping provides satisfactory alignment over Centrifuge classification. This filtering method was tuned to remove all hits from the negative controls but leave as many validated positive detection species reads as possible. It is therefore a heuristic method and can be tuned with greater power when more samples have been processed.

**Figure 3.**
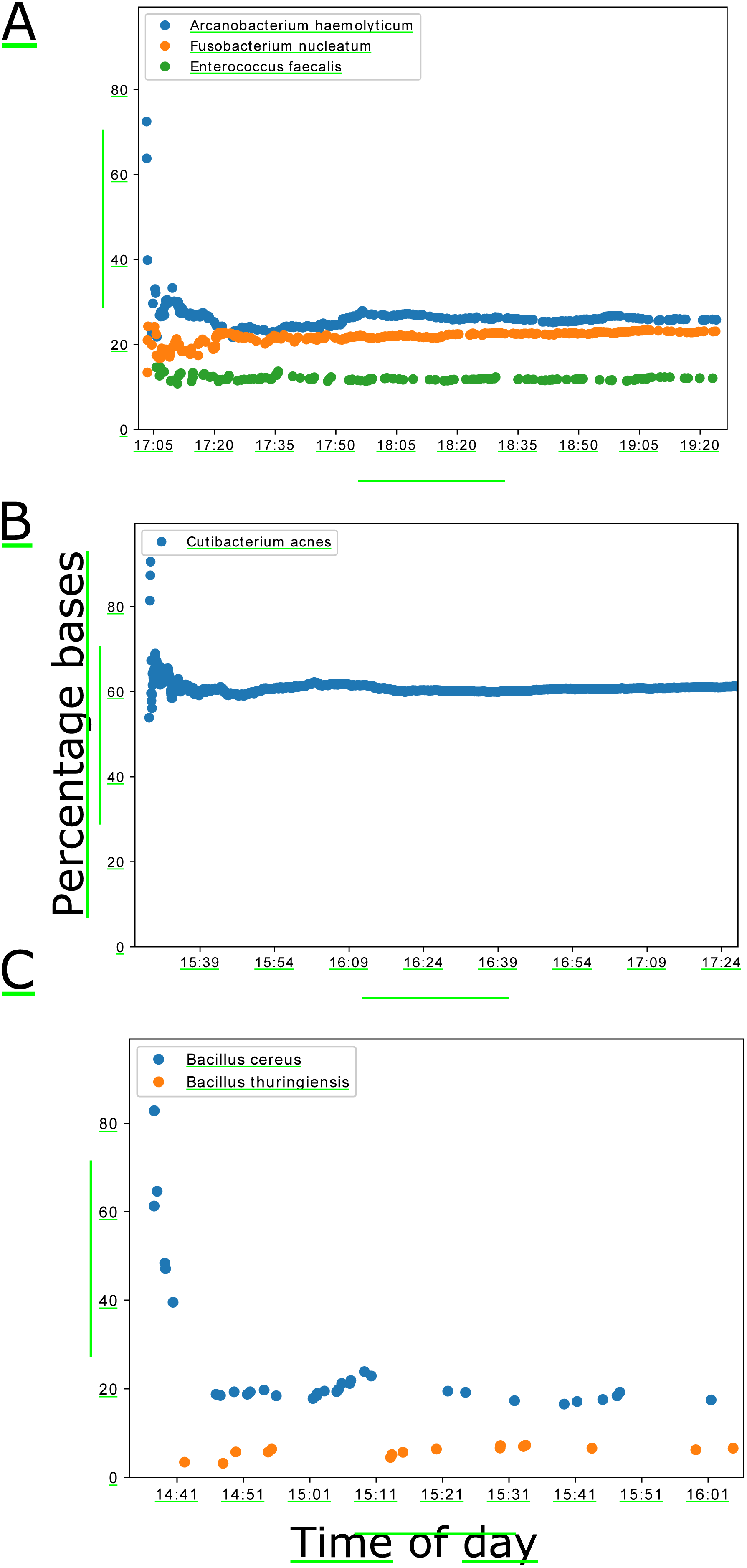
Percentage of mapped bases (minimap2) to total centrifuge classified bacterial bases over the first two hours of sequencing. As with Figure 2, each marker on the plots represents a new sequence classified. Times are on the day of sequencing. Three samples shown showcasing the best and worst performers. (A) Sample 354a containing three different species. (B) Sample 249a containing Cutibacterium acne. (C) Sample 352a containing two different Bacillus species.

The entire workflow was run in Nextflow [14] with the software contained inside a Singularity [15] image. This has enabled the entire pipeline to run on a distributed cluster (SLURM [24]) with the flexibility to run on other platforms. A SLURM cluster was setup and used to handle the high computational demands of basecalling with Albacore, with the remaining pipeline requiring less computer time to complete. The cluster setup was built from a head node and four worker nodes with a total of 21 worker cores. Centrifuge was only run on two of the nodes, each with at least 16gb of memory. The workflow can be run in real time and detect new fast5 files from a MinION sequencing run, process them and push the data to a MongoDB database for analysis.

## Results

### Sample composition after analysis

Nine samples previously sequenced with an Illumina MiSeq were sequenced using the Oxford Nanopore MinION platform. Seven samples were extracted from bacterial culture positive sonication fluid. The remaining two samples, extracted from culture negative sonication fluid, were used as negative controls. Between 0.2 and 2.8 gigabases were basecalled for each sequencing run, with read lengths averaging between 500 bp and 1.7 kb (**Error! Reference source not found**.).

The majority of classified reads were human, Table 1, with a range of 80% to 97% of bases in the sequenced culture positive samples coming from host contamination. Of the remaining non-host bases, a range of 0.04% to over 6% of bases were classified as bacterial by Centrifuge in the culture positive samples, Table 1.

**Table 1.** Oxford nanopore technologies MinION sequencing yields and basic details and breakdown of centrifuge classification. Bacteria, Human with a centrifuge score greater than 150, and total reads including unclassified reads. Samples 509a and 506a are culture negatives and used as negative controls. Results are after removing low complexity reads.

| Sample | Total |  | Mean | Median | Low complexity |  | Human |  | Bacteria |  |
| --- | --- | --- | --- | --- | --- | --- | --- | --- | --- | --- |
|  | Bases | Reads | read | read | bases | reads | bases | reads | bases | reads |
|  |  |  | length | length |  |  |  |  |  |  |
| 229a | 204,346,556 | 124,218 | 1,645.06 | 1,745 | 113,836 | 69 | 198,972,861 | 117,821 | 1,692,097 | 914 |
| 249a | 723,925,562 | 585,098 | 1,237.27 | 1,006 | 403,668 | 370 | 563,888,189 | 411,612 | 44,502,912 | 34,773 |
| 259a | 1,057,865,247 | 600,291 | 1,762.25 | 1,321 | 390,209 | 289 | 949,663,786 | 502,426 | 512,827 | 312 |
| 312a | 1,121,119,742 | 1,004,818 | 1,115.74 | 674 | 1,905,423 | 1,044 | 1,038,235,876 | 882,763 | 30,426,198 | 14,948 |
| 335a | 2,847,687,425 | 1,717,810 | 1,657.74 | 1,171 | 2,835,054 | 1,362 | 2,783,128,118 | 1,605,466 | 1,388,748 | 989 |
| 352a | 803,638,340 | 986,867 | 814.33 | 609 | 567,656 | 630 | 669,796,136 | 752,022 | 459,779 | 579 |
| 354a | 706,380,170 | 945,929 | 746.76 | 596 | 680,560 | 848 | 570,485,740 | 717,662 | 2,151,551 | 2,443 |
| 509a | 2,740,060,527 | 4,940,241 | 554.64 | 439 | 16,355,839 | 24,413 | 1,199,779,866 | 1,352,438 | 6,240,425 | 2,628 |
| 506a | 2,451,399,949 | 4,700,013 | 521.57 | 431 | 20,014,343 | 23,631 | 1,161,796,584 | 1,671,726 | 4,705,919 | 2,139 |

Our analysis workflow identified one or more bacterial species per sample, with the exception of the two culture negative samples, 509a and 506a (Table 2). One sample, 354a, was polymicrobial, with *Enterococcus faecalis, Arcanobacterium haemolyticum* and *Fusobacterium nucleatum* identified. Two species of the same genus, *Bacillus cereus* and *Bacillus thuringiensis,* were identified in sample 352a. All other samples had only a single bacterial species identified.

**Table 2.**
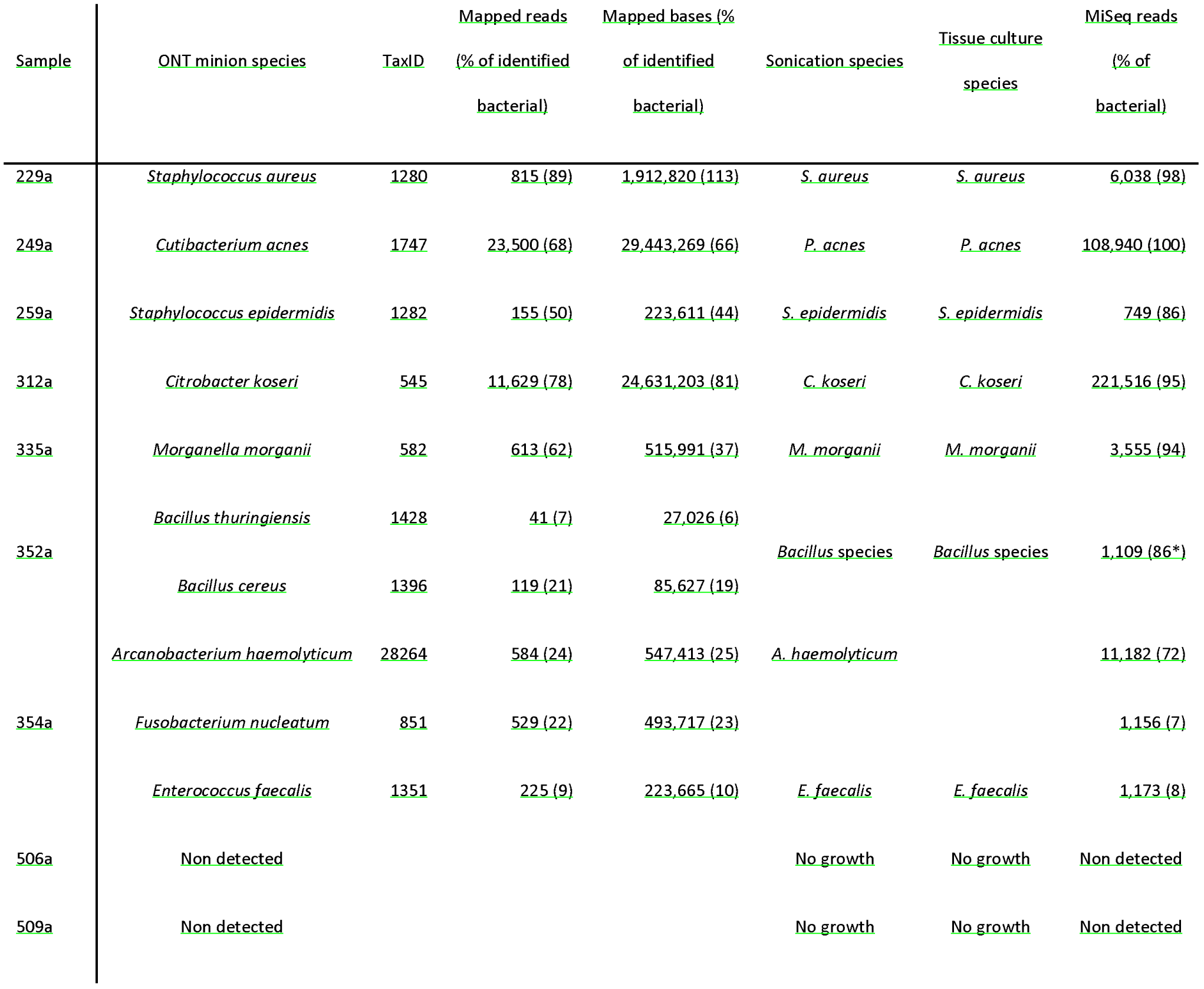
Species detected after read classification and reference genome alignment in CRuMPIT. Samples 509a and 506a are culture negatives and used as negative controls, no bacterial species were detected after filtering thresholds were used. Species detected from sonication fluid, tissue culture and MiSeq sequence analysis using Kraken. Adapted from [8]. (*) indicates % of bacterial reads taken from the Bacillus cereus group level (taxonomic id of 86661).

The results from ONT MinION sequencing correspond with previously published analysis of the same samples by conventional microbiology culture and metagenomic Illumina MiSeq sequencing, Table 1 [8]. A notable difference between the two molecular analyses can be seen in sample 352a, where ONT MinION sequencing enabled species level detection. The Illumina short read sequencing identified *Bacillus spp.* only (agreeing with the corresponding culture results) whereas ONT MinION sequencing identified two species from the *Bacillus cereus group: Bacillus cereus* and *Bacillus thuringiensis.* It is worth noting that speciation within the *Bacillus cerus group* is problematic as species within this group share a high level of genome sequence identity [25]. Further investigation would be required to determine whether both species are actually present in this sample.

Another difference observed between the two sequencing techniques is in sample 354a, and concerns the relative abundance of sequencing reads/bases for the multiple species classified in this polymicrobial sample. The Illumina MiSeq sequencing identified *A. haemolyticum* as the most abundant species, at 72% of bacterial reads, with F. *nucleatum* representing 7% of bacterial reads. However, ONT MinION sequencing classified very similar base numbers for both *F. nucleatum* and A *haemolyticum* (493,717 and 547,413 bases respectively) We speculated that this observed difference in proportions of reads for the *F. nucleatum* and *A. haemolyticum* was caused by platform sequencing bias, possibly as a result of variable genome GC content: The *A. haemolyticum* genome is 54% GC, compared to 27% for F. *nucleatum.* We used qPCR to test our hypothesis, and investigate which platform represents an estimate of genome abundance of these two species that is closest to the original DNA extract from sample 354a. qPCR results detected approximately equal copy numbers of both *A. haemolyticum* and F. *nucleatum* genomes in the original DNA extract, suggesting that ONT MinION sequencing has given a more accurate representation of species abundance in sample 354a, Table 4. However, standard deviations were high therefore further investigation will be needed to confirm this.

**Table 3.**
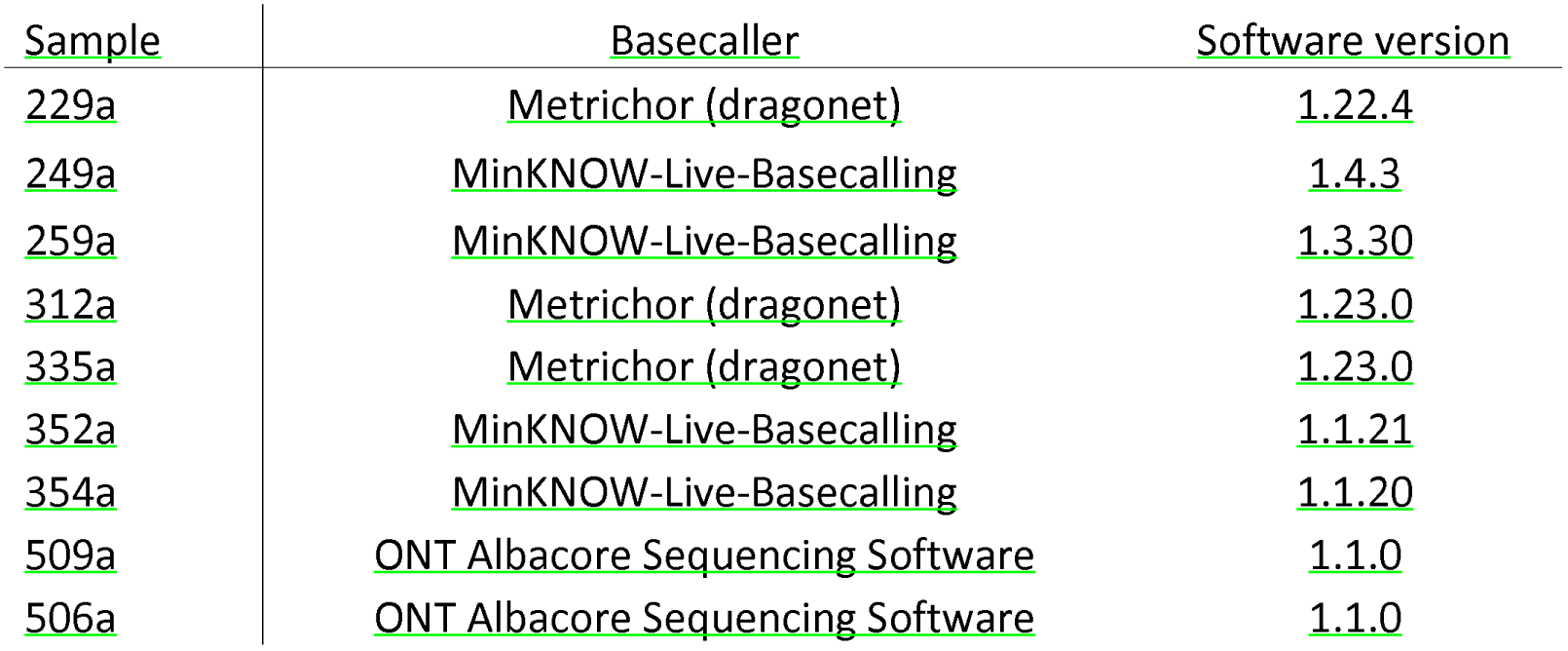
Nanopore basecallers and versions used for each sample.

| Sample | Basecaller | Software version |
| --- | --- | --- |
| 229a | Metrichor (dragonet) | 1.22.4 |
| 249a | MinKNOW-Live-Basecalling | 1.4.3 |
| 259a | MinKNOW-Live-Basecalling | 1.3.30 |
| 312a | Metrichor (dragonet) | 1.23.0 |
| 335a | Metrichor (dragonet) | 1.23.0 |
| 352a | MinKNOW-Live-Basecalling | 1.1.21 |
| 354a | MinKNOW-Live-Basecalling | 1.1.20 |
| 509a | ONT Albacore Sequencing Software | 1.1.0 |
| 506a | ONT Albacore Sequencing Software | 1.1.0 |

**Table 4.** qPCR results.

| Species | Std curve RSq | Efficiency | Replicate | CT | Copies | Average ± Std Dev |
| --- | --- | --- | --- | --- | --- | --- |
| *Arcanobacterium haemolyticum* | 0.991 | 89.20% | 1 | 29.12 | 2356 | 3214 ± 965 |
|  |  |  | 2 | 28.72 | 3028 |  |
|  |  |  | 3 | 28.19 | 4258 |  |
| *Fusobacterium nucleatum* | 0.999 | 86.00% | 1 | 28.93 | 4269 | 3421 ± 1304 |
|  |  |  | 2 | 30.22 | 1919 |  |
|  |  |  | 3 | 29.01 | 4075 |  |

### Real time analysis

Using the ONT MinION platform, it was possible to analyse sequences in real-time, and predict the species composition of culture positive samples minutes after data acquisition. Samples containing a larger yield of bacterial DNA, such as 354a and 249a, produced several hundred kilobases of sequences within the first two of hours, Figure 2A+B. Samples with lower yields, such as 352a, produced less sequence data, with several kilobases generated in the first two hours, Figure 2C. For all the species identified that passed the analysis thresholds, however, the sequences generated after data acquisition were consistent with the species identified by traditional culture methods and MiSeq sequencing, Figure 3. Each batch analysed within the Nextflow workflow took between four and fifteen minutes to process using a single core, depending on which node the job was submitted to, Supplemental figure 5. Therefore, real-time in this context needs to include this bioinformatics analysis time, the majority of which is basecalling.

## Discussion

Here we demonstrate proof-of-principle that long-read sequencing using the ONT MinION can detect bacterial infections from DNA extracted directly from sonication fluid samples, and potentially do so within minutes of starting sequencing. If DNA extraction techniques can be similarly optimised, these technologies have the potential to make intra-operative diagnosis of the causes of specific infections possible. This would allow both local and systemic antibiotics to be targeted to the causative organisms in prosthetic joint infections, starting at the time of surgery.

Analysis of the MinION data indicates concordance with the current gold standard laboratory culture and also Illumina short-read sequencing. In addition, we present a new analytical tool, CRuMPIT, which automates analysis of MinION data in this setting, and could be applied by other researchers and clinicians. By using negative controls we were able to determine signatures of background contamination - a challenge to diagnostic metagenomic interpretation [8, 26]. The thresholds and scores used within our bioinformatics workflow were determined after sequencing two negative controls that allow us to create heuristic thresholds to remove background sequences and false positives without masking the infection species. Future studies will involve sequencing more samples so refined threshold scores can be determined. This can be done as before with a Youden Index and J-statistic [8]. Sensitivity and specificity of the MinION cannot be determined from this study and therefore further, more extensive studies are required before use in a routine diagnostic microbiology laboratory can be recommended.

Although we were able to predict each species present within the sequenced samples, the vast majority of DNA sequenced was human, from host contamination, despite efforts to reduce this in the laboratory preparation. Depletion of host DNA contamination will facilitate greater pathogen genome sequencing coverage but this continues to present challenges as the numbers of bacterial cells in joint infections is low [5] in relation to human cells. Previous studies with ONT MinlON on direct clinical samples have used samples with relatively high concentrations of bacteria in urine [11] (compared to PJI samples) or moderate to high viral titres in blood [10]. The MinION has also been used for metagenomics in environmental samples [27]. However, reduction of human DNA could allow better genotyping, transmission analysis and antimicrobial resistance gene prediction as the proportion of bacterial DNA increases. Currently, this depends on laboratory development to reduce the number of human cells in samples rather than downstream bioinformatic analysis.

The sequencing yields here were low compared to other ONT MinION sequencing yields sequenced within the same lab (data not shown). DNA read lengths sequenced in this project are also relatively short, with the average under 1 kilobase, where mean read lengths can be expected over 10 kilobases with this method. This is likely due to the DNA extraction methods used, as they were optimised for MiSeq sequencing. However, of the four samples processed by PCR due to low DNA concentration, there was variation in read length and depth ranging from highest to lowest.

There are known biases for organisms associated with GC content in using PCR-based methods for sample preparation [28] and with Illumina metagenomic data [29]. We found some evidence that MinlON sequencing may better reflect the relative abundance of pathogen DNA in polymicrobial infections, as it appeared less prone to GC biases than Illumina MiSeq short-read sequencing.

Detection of the species was possible within minutes of the sequencing run starting, and this includes the time required to process the sequencing data, with basecalling being the biggest bottleneck. The fast5 file batch size has an effect on turnaround time and reducing batch sizes is preferable for longer reads that take more time to basecall. We have tested the pipeline on a SLURM cluster on the same network as the computers running the MinION sequencer, enabling us to scale to the rate of sequencing and basecall with greater throughput than we could with a single machine.

A limitation of this study was seen in runs where reads were live basecalled with the MinKNOW basecaller: the runs produced data too quickly for the system to keep up. Retrospective basecalling was not possible at the time and the skipped reads have since been discarded. Therefore, in future studies using Albacore, as is the case with the most recent two sequencing runs (506a and 509a), we expect the average DNA read lengths and yields to increase, which will aid species classification and potential *De novo* assemblies.

The ONT MinION sequencing process has undergone continual development with substantial improvements since this project began. Therefore, we have used three different basecallers, Metrichor, MinKNOW and Albacore, for converting the raw signal or event data to DNA sequences. It is possible to rebasecall some of this data, but as we no longer have access to some sample raw data files, we cannot rebasecall all the samples. Also, as this would not reflect the real-time analysis carried out, we have not rebasecalled all samples with the same software version. Future studies should continue to use the most accurate, current, and efficient basecaller for real-time analysis.

Although analysis of the sequencing is close to real-time, the DNA extraction and library preparation takes several hours. There are rapid library preparation kits available, however we feel the sequencing yield is currently too low for these to be a viable route to detection of pathogens directly from samples, particularly in samples with high host contamination. This project was a proof of concept, but to be cost effective in the future, multiplexed samples or reusable/washable flowcells may need to be employed.

## Conclusions

The study shows reliable detection of infection species composition in prosthetic joint infections using ONT MinION sequencing. This represents proof of concept for utilising real time ONT MinlON sequencing for PJI diagnostics. The speed of detection indicates that this technology has the potential to deliver results to the clinician in a timelier manner than traditional microbiological methods. Reduction of diagnostic time could have a significant positive influence on patient outcome, allowing prompt, targeted antimicrobial therapy.

The development of a reproducible workflow, as described in this study, has potential use for any clinical sample metagenomic ONT MinION sequencing, not just sonication fluids. The software used for analysis is provided [17] and can be installed and run locally or in a distributed cluster to scale with throughput.

---

Supplemental figure 1. Bases classified total or target over centrifuge score. Each sample has two lines of the same colour. The top line is total bacterial bases identified by centrifuge over the score threshold used. The second lower line is the validated detected species/infection for the sample (Target). As the score threshold increases, the number of total classified bases reduces at a great rate than the target bases, until a plateau and diminishing returns at approximately 150.

Supplemental figure 2. Each species identified by centrifuge showing total bases over number of reads as proportions of total bacterial bases and total bacterial reads respectively. Species detections below the 0.1 proportion (i.e. less than 10%) of bases threshold are dots and species detections above the 0.1 proportion threshold are crosses. Culture negative controls are red and Culture negative positive samples are blue.

Supplemental figure 3. Indiscriminate(indis) read and discriminate(dis) mapping qualities. Quality scores taken from mapping all reads to a reference with minimap2. Discriminate scores are from reads that have passed through the pipeline filtering thresholds and are determined to be reads specific to the reference. The indiscriminate are other reads that were likely to be host and/or contamination.

Supplemental figure 4. Each species identified by minimap2 mapping showing total bases over number of reads as proportions of total bacterial bases (centrifuge) and total bacterial reads (centrifuge) respectively. Species detections below the 0.1 proportion (i.e. less than 1%) of bases threshold are dots and species detections above the 0.01 proportion threshold are crosses. Culture negative controls are red and Culture negative positive samples are blue. Shows shortened axis of below threshold hits.

Supplemental figure 5. Batch job duration times in minutes sample report taken from Nextflow. Using sample 354a as a representative for the bioinformatic analysis. Batches were run over a heterogeneous SLURM cluster with variable node CPU speeds affecting Albacore performance.

## Declarations

### Ethics approval and consent to participate

For this study, no ethical review was required, because the study was a laboratory method development study focusing on bacterial DNA extracted from discarded samples identified only by laboratory numbers, with no personal or identifiable data. Sequencing reads identified as human on the basis of Centrifuge were counted and immediately discarded.

### Consent for publication

Not applicable.

### Availability of data and material

The raw FASTQ sequencing data from this publication are archived to EBI ENA with the bioproject identifier, PRJEB23460, and available to download. We hope to make the constructed centrifuge index available to download when we find a host for it and future iterations, until then please contact the corresponding author for access.

## Authors’ contributions

NDS performed bioinformatic analysis including creating CRuMPIT and wrote the manuscript. TLS performed all molecular laboratory work including DNA extraction, library preparation, sequencing run and qPCR. TLS,DF,TEAP and DWE helped prepare manuscript. JS helped setup computational infrastructure. TEAP, DWE and DWC contributed intellectually to the direction of the project and helped finish manuscript. BLA,AJB,MAM,SO and AT provided and processed biological samples.

## Acknowledgements

The authors thank the microbiology laboratory staff of the John Radcliffe Hospital, Oxford University Hospitals NHS Foundation Trust, for providing assistance with sample collection and processing. The views expressed are those of the author(s) and not necessarily those of the NHS, the NIHR or the Department of Health.

## References

1. Huotari K, Peltola M, Jamsen E. The incidence of late prosthetic joint infections: a registry-based study of 112,708 primary hip and knee replacements. Acta Orthop. 2015;86:321–5. doi:10.3109/17453674.2015.1035173.

2. Lenguerrand E, Whitehouse MR, Beswick AD, Jones SA, Porter ML, Blom AW. Revision for prosthetic joint infection following hip arthroplasty: Evidence from the National Joint Registry. Bone J. Res. 2017;6:391–8. doi:10.1302/2046-3758.66.BJR-2017-0003.R1.

3. Lenguerrand E, Whitehouse MR, Beswick AD, Toms AD, Porter ML, Blom AW, et al. Description of the rates, trends and surgical burden associated with revision for prosthetic joint infection following primary and revision knee replacements in England and Wales: an analysis of the National Joint Registry for England, Wales, Northern Ire. BMJ Open. 2017;7:e014056. doi:10.1136/bmjopen-2016-014056.

4. Rochford ET, Richards RG, Moriarty TF. Influence of material on the development of device-associated infections. Clin Microbiol Infect. 2012;18:1162–7. doi:10.1111/j.1469-0691.2012.04002.x.

5. Atkins BL, Athanasou N, Deeks JJ, Crook DW, Simpson H, Peto TE, et al. Prospective evaluation of criteria for microbiological diagnosis of prosthetic-joint infection at revision arthroplasty. The OSIRIS Collaborative Study Group. J Clin Microbiol. 1998;36:2932–9. https://www.ncbi.nlm.nih.gov/pubmed/9738046.

6. Osmon DR, Berbari EF, Berendt AR, Lew D, Zimmerli W, Steckelberg JM, et al. Diagnosis and management of prosthetic joint infection: clinical practice guidelines by the Infectious Diseases Society of America. Clin Infect Dis. 2013;56:e1–25. doi:10.1093/cid/cis803.

7. Bejon P, Berendt A, Atkins BL, Green N, Parry H, Masters S, et al. Two-stage revision for prosthetic joint infection: predictors of outcome and the role of reimplantation microbiology. J Antimicrob Chemother. 2010;65:569–75. doi:10.1093/jac/dkp469.

8. Street TL, Sanderson ND, Atkins BL, Brent AJ, Cole K, Foster D, et al. Molecular diagnosis of orthopaedic device infection direct from sonication fluid by metagenomic sequencing. J Clin Microbiol. 2017; August:JCM.00462–17. doi:10.1128/JCM.00462-17.

9. Ruppe E, Lazarevic V, Girard M, Mouton W, Ferry T, Laurent F, et al. Clinical metagenomics of bone and joint infections: a proof of concept study. Sci Rep. 2017;7:7718. doi:10.1038/s41598-017- 07546-5.

10. Greninger AL, Naccache SN, Federman S, Yu G, Mbala P, Bres V, et al. Rapid metagenomic identification of viral pathogens in clinical samples by real-time nanopore sequencing analysis. Genome Med. 2015;7:99. doi:10.1186/s13073-015-0220-9.

11. Schmidt K, Mwaigwisya S, Crossman LC, Doumith M, Munroe D, Pires C, et al. Identification of bacterial pathogens and antimicrobial resistance directly from clinical urines by nanopore-based metagenomic sequencing. J Antimicrob Chemother. 2017;72:104–14. doi:10.1093/jac/dkw397.

12. Mitsuhashi S, Kryukov K, Nakagawa S, Takeuchi JS, Shiraishi Y, Asano K, et al. A portable system for rapid bacterial composition analysis using a nanopore-based sequencer and laptop computer. Sci Rep. 2017;7:5657. doi:10.1038/s41598-017-05772-5.

13. Hassan AA, ülbegi-Mohyla H, Kanbar T, Alber J, Lämmler C, Abdulmawjood A, et al. Phenotypic and genotypic characterization of arcanobacterium haemolyticum isolates from infections of horses. J Clin Microbiol. 2009;47:124–8.

14. Di Tommaso P, Chatzou M, Floden EW, Barja PP, Palumbo E, Notredame C. Nextflow enables reproducible computational workflows. Nat Biotechnol. 2017;35:316–9. doi:10.1038/nbt.3820.

15. Kurtzer GM, Sochat V, Bauer MW. Singularity: Scientific containers for mobility of compute. PLoS One. 2017;12:1–20.

16. What is docker. 2017. https://www.docker.com/what-docker.

17. Sanderson ND. Clinincal Real-time Metagenomics Pathogen Identification Test (CRuMPIT).https://gitlab.com/ModernisingMedicalMicrobiology/CRuMPIT.

18. Wick RR, Judd LM, Holt KE. Comparison Of Oxford Nanopore Basecalling Tools. 2017. doi:10.5281/ZENODO.1043612.

19. Sanderson ND. fast5watcher.py. 2017. https://github.com/nick297/fast5_scripts.

20. Schmieder R, Edwards R. Quality control and preprocessing of metagenomic datasets. Bioinformatics. 2011;27:863–4.

21. Kim D, Song L, Breitwieser FP, Salzberg SL. Centrifuge: Rapid and sensitive classification of metagenomic sequences. Genome Res. 2016;26:1721–9.

22. Wood DE, Salzberg SL. Kraken: ultrafast metagenomic sequence classification using exact alignments. Genome Biol. 2014;15:R46. doi:10.1186/gb-2014-15-3-r46.

23. Li H. Minimap2: fast pairwise alignment for long nucleotide sequences. 2017;:2–5. doi:10.1101/169557.

24. Jette MA, Yoo AB, Grondona M. SLURM: Simple Linux Utility for Resource Management. In: In Lecture Notes in Computer Science: Proceedings of Job Scheduling Strategies for Parallel Processing (JSSPP) 2003. Springer-Verlag; 2002. p. 44–60.

25. Helgason E, økstad OA, Caugant DA, Johansen HA, Fouet A, Mock M, et al. Bacillus anthracis, Bacillus cereus, and bacillus thuringiensis-One species on the basis of genetic evidence. Appl Environ Microbiol. 2000;66:2627–30.

26. Kearney MF, Spindler J, Wiegand A, Shao W, Anderson EM, Maldarelli F, et al. Multiple sources of contamination in samples from patients reported to have XMRV infection. PLoS One. 2012;7.

27. Brown BL, Watson M, Minot SS, Rivera MC, Franklin RB. MinION nanopore sequencing of environmental metagenomes: A synthetic approach. Gigascience. 2017;6:1–10.

28. Jones MB, Highlander SK, Anderson EL, Li W, Dayrit M, Klitgord N, et al. Library preparation methodology can influence genomic and functional predictions in human microbiome research. Proc Natl Acad Sci. 2015;112:14024–9. doi:10.1073/pnas.1519288112.

29. Schirmer M, D’Amore R, Ijaz UZ, Hall N, Quince C. Illumina error profiles: resolving fine-scale variation in metagenomic sequencing data. BMC Bioinformatics. 2016;17:125. doi:10.1186/s12859-016-0976-y.

